# Opportunities and challenges for improving a Colombian public research program in plant breeding and plant genetic resources lead by Agrosavia

**DOI:** 10.1101/2020.09.21.305961

**Authors:** Carlos H. Galeano, Katherine Tehelen, Hugo R. Jiménez, Carolina González, Ivania Cerón-Souza

**Author notes:** **Author contributions:** Carlos H. Galeano, Hugo Jiménez, and Carolina González designed the study. Carlos H. Galeano, Katherine Tehelen, and Ivania Cerón-Souza performed the data analyses and interpretation. All authors wrote the manuscript and approved the final version of the manuscript for publication.

## Abstract

Agrosavia (Corporación Colombiana de Investigación Agropecuaria) is the Colombian state institution in charge of the agricultural research at the national level, including plant breeding. Since 2014, Agrosavia started to increase its research staff and has reset the leadership of public research to solve the needs of the agricultural sector population, focusing on small producers. However, the current team working on plant breeding and plant genetic resources are facing some challenges associated with generation gaps and the lack of a collaborative working plan for the next years. To identify the opportunities and actions in this research area, we surveyed all the 52 researchers working in Agrosavia in this area in 2017. We analyzed the opinions of researchers to detect the strengths and weaknesses of the program using a sentiment score. We also examined the networking to test both how consolidated the group is and if among top leaders are gender parity and also have a higher academic degree. Results showed that there is a mixed community of old and new researchers with clear gender bias in the proportion of male-female. Within the network, the interactions are weak, with several subgroups where the top-ten of both central leaders and the most influencer are frequently males with mostly an M.Sc. degree but with significant experience in the area. Researchers have an interest in 31 crops. From them, 26 are in the national germplasm bank, but this bank is not the primary source for their breeding programs. The top-five of plants with increasing interest are corn, cocoa tree, golden berries, oil palm, and sugarcane. Researchers also want to establish collaborations with 54 different institutions, where the *Universidad Nacional de Colombia*, which is the top public university in the country, is on the head. Researchers also perceived weaknesses in the marker-assisted selection, experimental design, and participatory plant breeding, but those criticisms have a positive sentiment score average of 1.55 (0.3 SE) across 31 texts analyzed. Based on all results, we identified five critical strategic principles to improve the plant-breeding research program. They include a gender diversity policy to hire new researchers strategically to reduce the gender gap and strength the generational shift. Better collaboration between the national germplasm bank and plant breeding research. A coordinate plan where the studies focus on food security crops that the government supports independently of market trends. And finally, adequate spaces for the project’s design and training programs. Hence, we recommend the creation of a consultant group to implement these policies progressively in the next years.

## Introduction

Plant breeding is a field where many complementary disciplines converge to generate more productive and better-adapted cultivars that respond to the challenges of world agriculture. Traditionally, plant breeding depends significantly on the selection of plants based on the evaluation of target agronomic traits, increasing the frequency of favorable alleles. The primary outcome of a plant breeding program is to increase crop productivity (i.e., yield, resistance to diseases and pests) and specific crop adaptation (i.e., tolerance to heat, frost, drought or salinity). Moreover, other desirable traits for plant breeding are the development of new processing alternatives (i.e., culinary quality, milling, fermentation, biofuel production, postharvest storage, and shelf life) and consumer acceptance (i.e., flavor, protein content, oil profile, fiber quality, nutritional value) [1].

Currently, the world requires more productive cultivars to provide enough food for the constantly growing human population, which will reach an extraordinary number of 9,000 million people by 2050. Therefore, world food production must increase from 25% to 70% compared with 2014 [2]. Moreover, due to global warming, there is an urgent need to develop varieties with high nutritional value and adaptation to stresses such as droughts, floods, frosts, and certain soils, but also to pests and diseases ([3,4]. In this context, new methodologies and tools based on genomics are helping in the generation of more accurate data for better characterization of the genetic diversity of crops and the association of critical phenotypic traits with genome regions [1,5].

In Colombia, research on plant breeding began in the 50s, directed by DIA (*Departamento de Investigaciones Agrarias*). At that time, a large group of experts developed the first successful plant varieties in different crops such as corn and potato (i.e., Diacol H104 and Diacol Capiro, respectively). During this period, DIA becomes ICA (*Instituto Colombiano Agropecuario*), starting an essential collaboration with the Rockefeller Foundation. In this period, research was focused actively in areas such as plant breeding, plant pathology, and plant physiology. A large group of researchers from ICA was granted with fellowships to carry out M.Sc. and Ph.D. studies, as well as research training outside the country. Besides, ICA led the creation and consolidation of the National Germplasm Bank of Colombia for agricultural purposes and also several genetic improvement programs with a multidisciplinary research scope [6]. Both government initiatives contributed to establishing the first national master’s degree program in genetics and the first breeding program in collaboration between ICA and *Universidad Nacional de Colombia*. During the next 30 years, ICA produced and released a large number of plant varieties, mainly corn, wheat, barley, oat, rice, and potato, which generated high impact and benefits for producers in the country. In the early 90s, the government created Agrosavia (previously known as Corpoica) as a national research institution in agriculture, leaving ICA only in charge of the phytosanitary control, inspection, and surveillance functions. Unfortunately, at the end of the 90s, government priorities changed, and the research budget for agricultural research, including plant breeding programs, was severely affected. Due to these unfortunate decisions, the national plant breeding research goal shifted to carry out field assessment of plant varieties developed by International Centers such as CIAT (International Center for Tropical Agriculture), CIMMYT (International Maize and Wheat Improvement Center) and CIP (International Potato Center).

From 2014, the national government, through Law Project No. 1731, approved by the Colombian Congress, promised to assign an annual budget to support and promote the agricultural research programs in Agrosavia. Since then, Agrosavia has increased its research capacity progressively through the incorporation of new research staff as well as new infrastructure. Currently, the central core of researchers working in plant breeding and plant genetic areas are young Ph.D. researchers trained in the use of both conventional and cutting-edge methodologies and tools for the selection and development of new plant varieties. However, Agrosavia is still working on different strategies to accomplish its objectives. From 2014 until 2016, the main focus was the creation of research networks based on national agricultural production chains. Then, from 2017 until today, Agrosavia has been working to consolidate disciplinary groups [7]. As part of this strategy, this study aims at characterizing the challenges and future opportunities for the current plant breeding and plant genetic resources (PB&PGR) group established in Agrosavia. Based on the results of this study, we identified the most important crops to future projects, the external institutions better to collaborate, and five critical strategic principles to strengthen a public plant breeding research program in Colombia lead by Agrosavia.

## Materials and methods

In February 2017, we surveyed the total of the 52 researchers from Agrosavia that, in that year, were working on PB&PRG projects. We designed an online Google questionnaire divided into three sections: (1) The group description, (2) Challenges and opportunities in the PB&PGR area, and (3) the construction of a social network based on the links within the PB&PRG group and externals (i.e., from other disciplines in Agrosavia and outside institutions) in six different scientific activities during the last ten years.

We performed all the statistical analyses and visualization using different R-packages under R Studio 1.2.5001. For section (1) and (2), we used Plotrix 3.7-4 [8] and GGPLOT2 3.3.1 [9] to compare age, gender identity, highest academic degree, years of experience, and rank category among the 52 researchers from the PB&PGR group in Agrosavia. Moreover, we identified and counted the crops that researchers worked in the last ten years and compared them with the list of plants that researchers want to continue working in the future and separate them into three categories, such as increasing, equal or decreasing interest among the 52 researchers. Furthermore, we counted how many times the researches have been using the National Germplasm Bank in their projects and the name of the external institutions that researchers consider key to working collaboratively in the future. Also, we asked the 52 researchers to agree or disagree with ten questions associated with identifying the strengths and weaknesses across different activities and skills in the area. The questions were: (1) Should PB&PGR group participates in advising and training students and young professionals?, (2) should PB&PGR group open spaces for journal clubs and research discussion? (3) Have students and young professionals a good knowledge of quantitative genetics? (4) Have students and young professionals a good understanding of fieldwork? (5) Have PB&PGR researchers a good understanding of MAS (Marker-Assisted Selection)? (6) Have PB&PGR researchers a good knowledge of experimental design? (7) Have PB&PGR researchers a good understanding of G (Genetic) x E (Environment) models? (8) Have PB&PGR researchers a good understanding of quantitative genetics? (9) Have PB&PGR researchers a good knowledge of molecular plant breeding? And (10) have PB&PGR researchers a good understanding of participatory plant breeding?

Finally, section (2) had an open question asking for comments or suggestions to improve the PB&PGR group in the future. Because we got 31 different answers, we made a corpus text mining analysis that included a sentiment analysis and a word cloud. The first step was to translate the 31 texts from Spanish to English using the free software DeepL (https://www.deepl.com/translator) and reviewed by hand securing the automatic translation was the closest possible to the original meaning. Then, we cleaned the corpus under the R-function GSUB and the R-package TM 0.7-7 to remove punctuations, numbers, empty spaces, stopwords, and a list of words associated with the workplace and research area (i.e., Agrosavia, plant breeding, plant genetic resources, etc.). Finally, using the R-package STRINGR 1.4.0, we broke each text into its characters to compare them with a positive and a negative list of words [10,11]. Both the positive and negative words identified weight one. Thus, we generated a sentiment score for each text subtracting the number of negative words from the number of positive words count. If the score was positive, the opinion has a positive sentiment. But, if the score was negative, the sense of the text is negative. We calculated the average and the standard error of the sentiment score across the 31 opinions. Moreover, we generated a word cloud based on the 30 most frequent words across the corpus under the R-package WORDCLOUD 2.6.

For section (3), we used the R-package IGRAPH 1.0.1 [12] to construct a two-way social network for analysis that consist in 48/52 researchers from the PB&PGR group that specified 812 interactions within the group but also with 116 external researchers (i.e., either with other disciplines within Agrosavia and outside institutions). We asked for six class of interactions among researchers, such as collaboration on projects in the PB&PGR area, discussion about new advances in the PB&PGR area, germplasm interchange or request, preparation of scientific papers and grey literature, registration of new varieties and technical advising. For each class, we compared the number of interactions within researchers from the PB&PGR group vs. between PB&PGR researchers and external.

In the network, we calculated and ranked the degree and the influence for each one of the researchers (i.e., vertex). The degree is a measurement of the number of adjacent interactions (i.e., edges). We associated the top-ten of the degree score, with the highest academic degree of the PB&PGR researchers to test the hypothesis that as a higher educational degree, higher interactions. Moreover, we measured the influence of each researcher based on two related statistics, the hub, and the authority scores. The hub score measures the interactions that a researcher has with different relevant influential researchers (i.e., authorities). Thus, the hub researchers are those that work together with trustworthy researchers on a common topic.

In comparison, the authority score measures the number of links of interactions that a researcher earned. Therefore, an authority researcher is one that should have relevant information on the field and, thus, received more links from other researchers. These two scores reinforce mutually because a good hub hint at many competent authorities; and, a proper authority is pointed to by many good hubs [13]. We associated the top-ten of both hub and authority scores with the gender information of PB&PGR researchers to test the hypothesis that researchers’ influence is independent of gender.

For the overall network, we calculated four descriptive statistics (i.e., the width, the edge density, the average distance, and the transitivity) to diagnose how researchers are working together within and outside the PB&PGR group. The diameter, the density, and the mean distance indicate together how connected the network is. In other words, they measure the ability of information to run through the system. Thus, for a tied network (i.e., with fewer intermediates across interactions and higher direct connection), we should expect lower diameter, higher density, and smaller mean distance. Likewise, the transitivity measures the local-scale-structure of the network. Thus, weak transitivity suggests that interactions occur in clusters loosely connected. In comparison, high transitivity indicates a well-consolidated system without a chance to identify discrete internal subgroups [14].

The methodology of this study was revised and approved by the biotetichs committee of Agrosavia. All researchers surveyed were verbally notified about the purpose of the study and voluntarily accepted to participate. Moreover, all the surveyed researchers knew and discussed the results during the second workshop of plant breeding and plant genetic resources celebrated in The Tibaitata Researc Center of Agrosavia during March 22-24, 2017.

## Results

### The group description

We found an apparent gender disparity among researchers working on PB&PGR within Agrosavia. The 71% (n=37) are male, and 29% (n=15) are female, and none declared a gender identity outside the binary male-female. The 40% (n=21) of both male and female researchers are between 31-40 years old. Male researchers are present in all age ranges, but female researchers are more frequent at younger intervals of age. There are only two female researchers in the age group of 51-70, and none are older than 60 years (Fig.1A).

Moreover, male researchers are present in all Agrosavia’S rank and all ranges of ages of experience within the PB&PGR group. In comparison, female researchers are only distributed in five of the eight categories, and with mostly 1-10 years of experience. However, no female researcher has more than 30 years of experience (Fig. 1B). Concerning the education background, the 52% (n=27) are M.Sc. researchers (20 male and seven female), and 27% (n=14) are Ph.D. researchers (seven male and seven female) (Fig. 1B). Considering lower academic degrees, both men and women studied only in Colombia. Nevertheless, there is a gender imbalance associated with studies abroad. From 37 male researchers, 14 (nine M.Sc. and five Ph.D.) studied abroad. In contrast, from 15 female researchers, four obtained their Ph.D. degree abroad (Fig. 1C).

**Figure 1.**
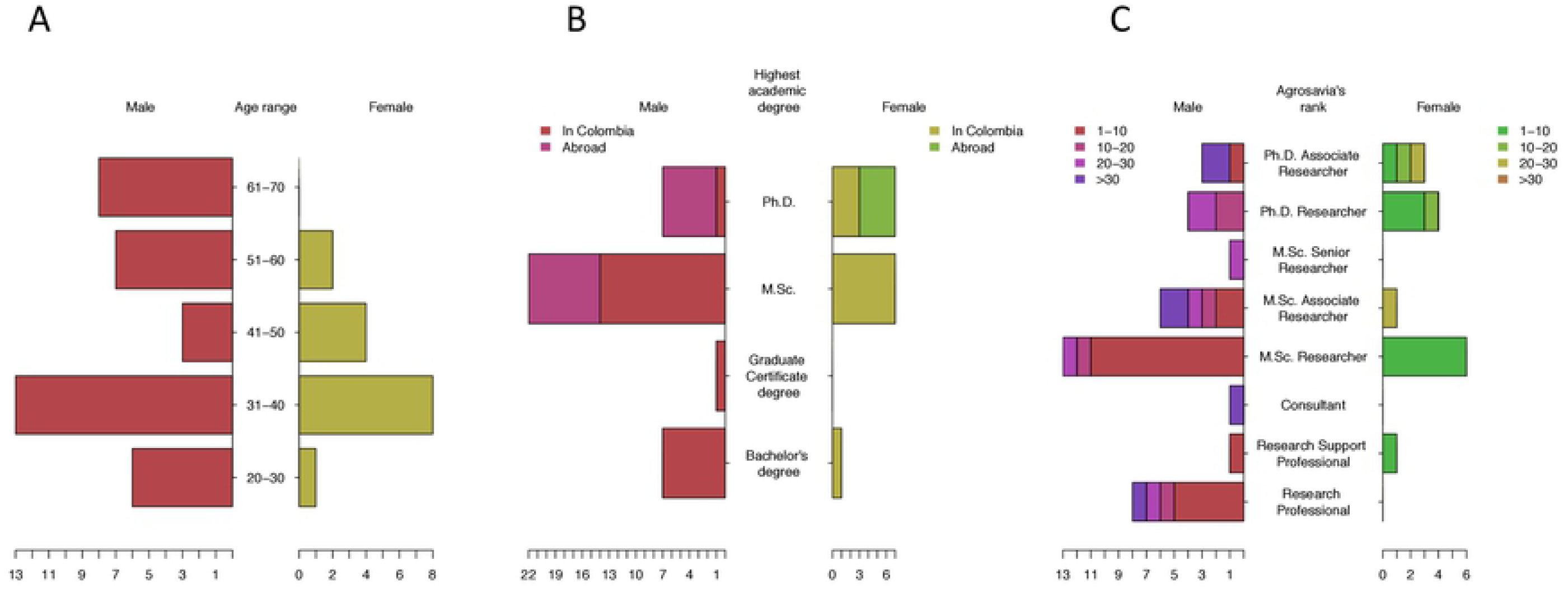
The characterization of a population of 52 researchers working in Agrosavia that participated in this study counting the number of researchers by (A) age range, (B) rank category in Agrosavia, and (C) the highest academic degree obtained in Colombia and abroad separated by gender. The horizontal scales represent the number of researchers, males at the left, and females at the right.

### Challenges and opportunities in the PB&PGR area

Researchers from Agrosavia have a broad spectrum of investigation experience in at least 31 crops. From this list, 19 species showed an increased research interest (i.e., more researchers interested in future projects than currently working on them), three species with equal importance between past and future research plans and nine species with decreased interest (i.e., few researchers interested in future projects than currently working on them) (Fig. 2). Although the National Germplasm bank conserves 26 of the 31 species from the list, the survey showed that this germplasm bank, also administered by Agrosavia, is not the primary focus of pre-breeding and breeding programs. The 37% (N=19) of the researchers have never used the germplasm bank, and 40% (N=21) have used once or twice in the last ten years (Fig. 3). Finally, researchers working on PB&PGR are interested in establishing collaborations with different 54 universities and research institutions inside and outside of Colombia in the next years. The most relevant was the *Universidad Nacional de Colombia*, with 23 researchers interested in collaborating. Moreover, 29 researchers want to establish partnerships with three of the CGIAR (Consultative Group for International Agricultural Research) bases within and outside of Colombia, such as CIAT (Colombia), CYMMIT (Mexico), and CIP (Peru). Finally, other research institutions with at least six researchers looking for collaborations are Embrapa (Brazil), CIRAD (France), and Cenicaña (Colombia) (Fig. 4).

**Figure 2.**
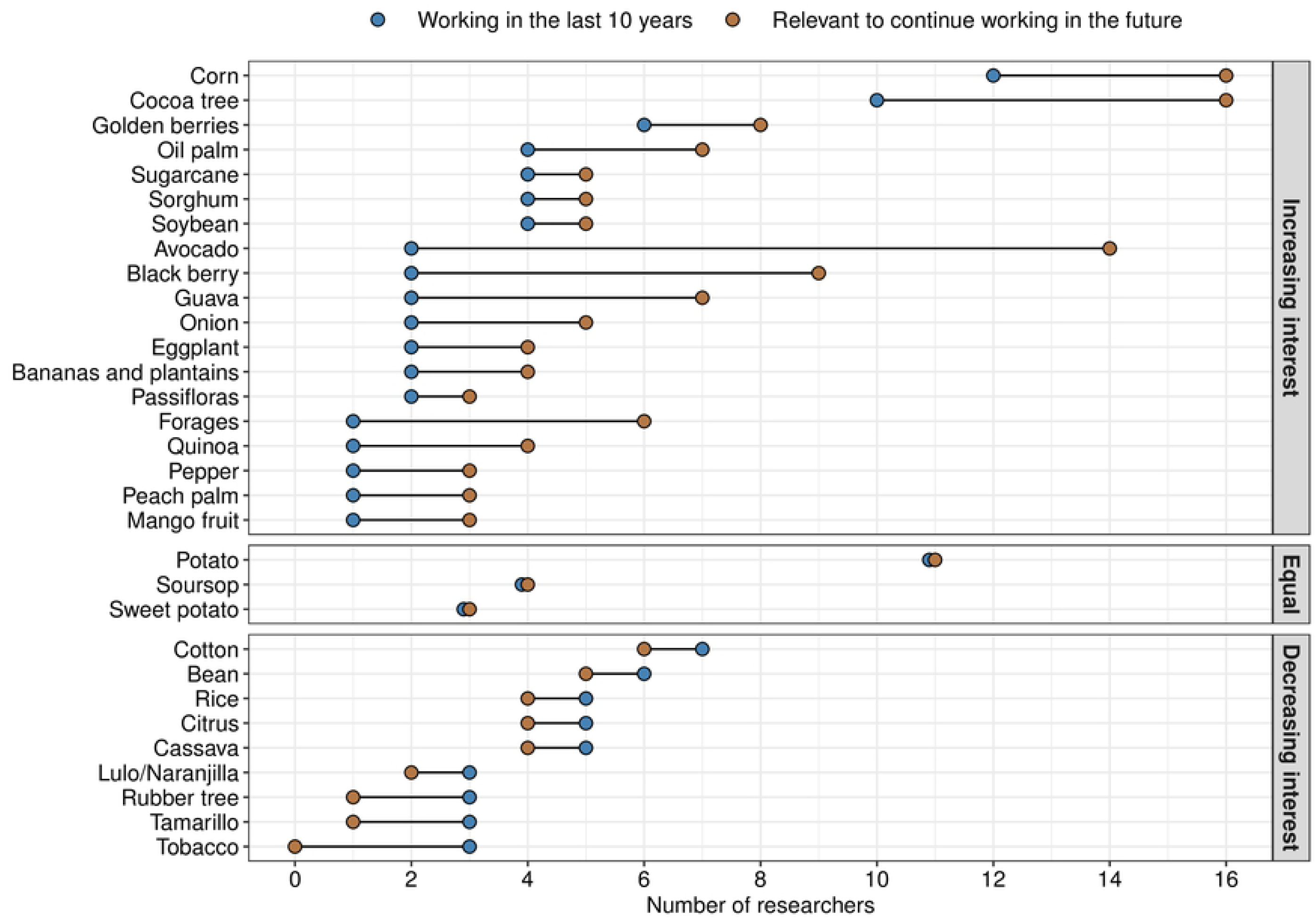
List of crops indicating in each line, the number of researchers in Agrosavia that work or have worked with these in the last ten years (blue dots), and the number of researchers that consider relevant to continue working with these in the future (yellow dots). There are three groups according to the patterns between past and future research interest, thus: crops with increasing interest, crops with similar interest in the historic vs. the future, and crops with decreasing interest in the future.

**Figure 3.**
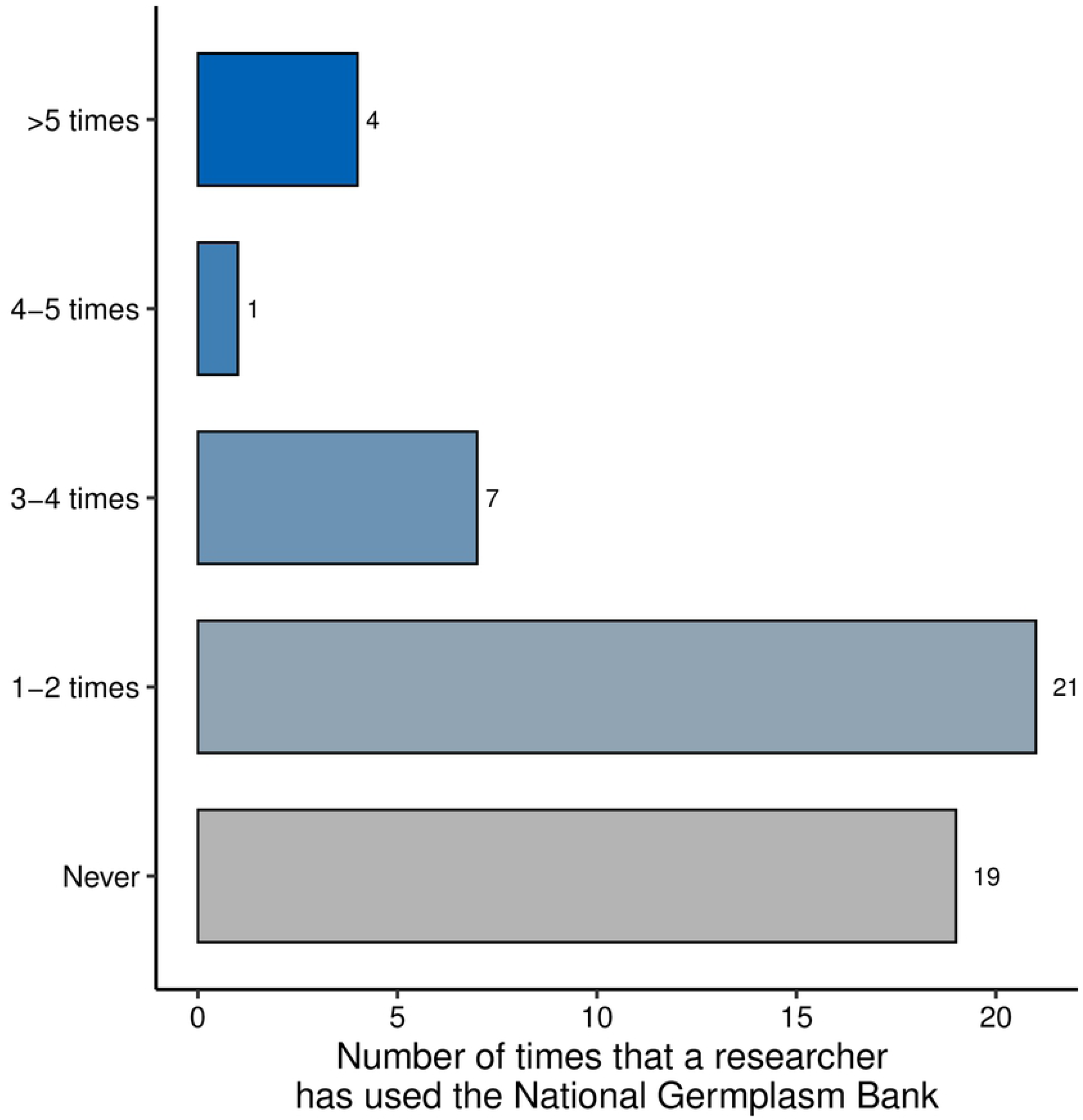
The frequency that researchers working on plant breeding and plant genetic resources from Agrosavia have requested germplasm from the National Plant Germplasm Bank of Colombia in the last ten years. The color of the bars is increasing from the five categories analyzed, from never to higher than five times.

**Figure 4.**
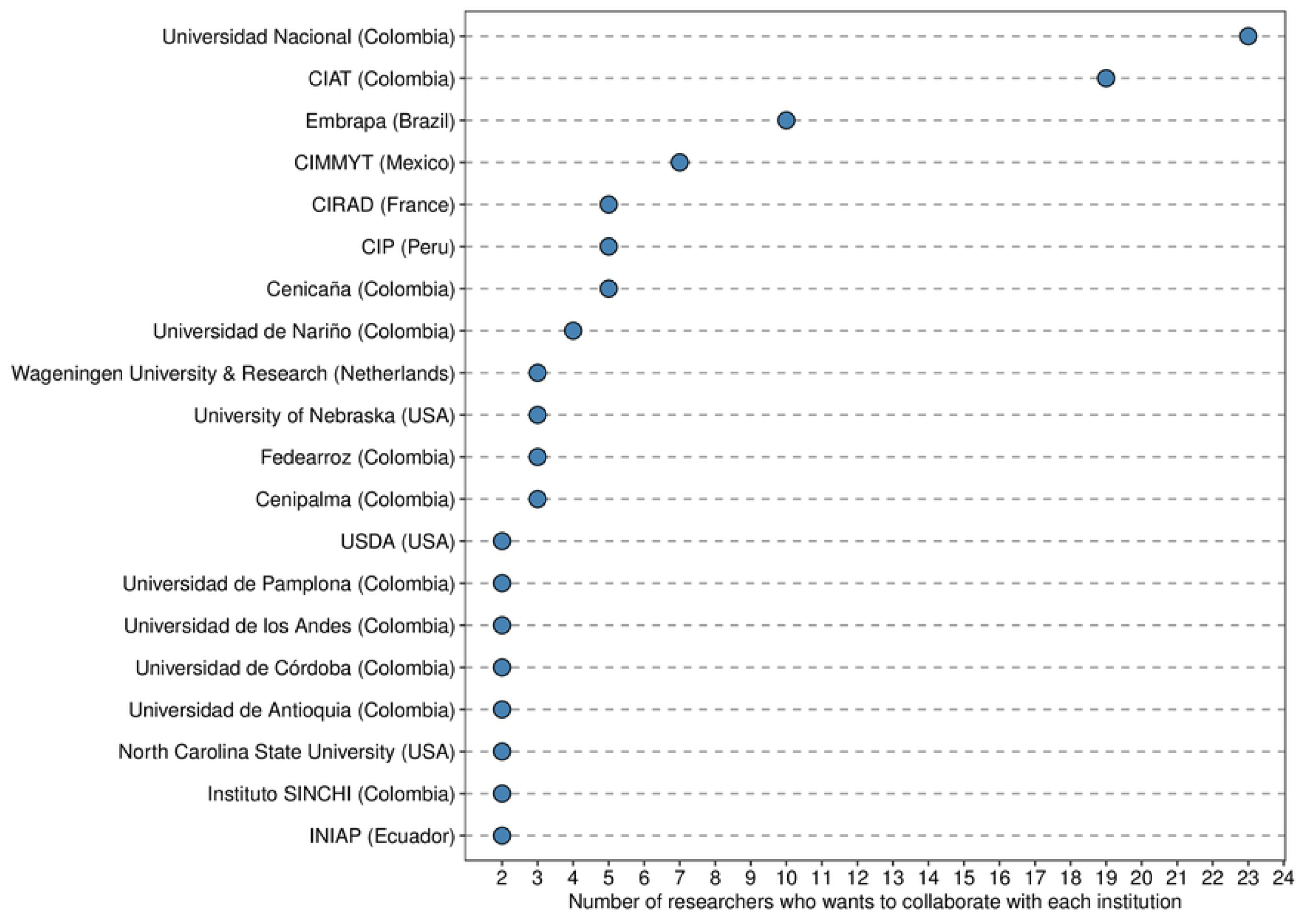
The number of researchers working on plant breeding and plant genetic resources from Agrosavia interested in starting collaborative breeding initiatives with national and international research institutions and universities. In parenthesis is the country of origin of each institution.

We found that researchers perceived weaknesses in fieldwork, Marker Assisted Selection (MAS), experimental design, and participatory plant breeding skills. Moreover, the researchers feel that they should contribute more actively to advise and train students and young professionals, as well as leading in journal clubs and research discussion spaces (Fig. 5). The opinion of 31 researchers about how to improve the group agreed in three main points: (1) work as a network where the different PB&PGR researchers benefit from the knowledge of others members of the group in several areas and disciplines, (2) work jointly with other researchers that are strong in other subjects such as plant physiology and plant phytopathology and (3) improve our impact by focusing on very few strategic crops but developing a multidisciplinary long-term research program. The top-five of the most used words include “programs, network, knowledge, improvement, and researchers” (Fig.6). The opinions and the most frequent words enclosed were purposeful, with a positive sentiment score average of 1.55 (0.3 SE) across 31 texts analyzed.

**Figure 5.**
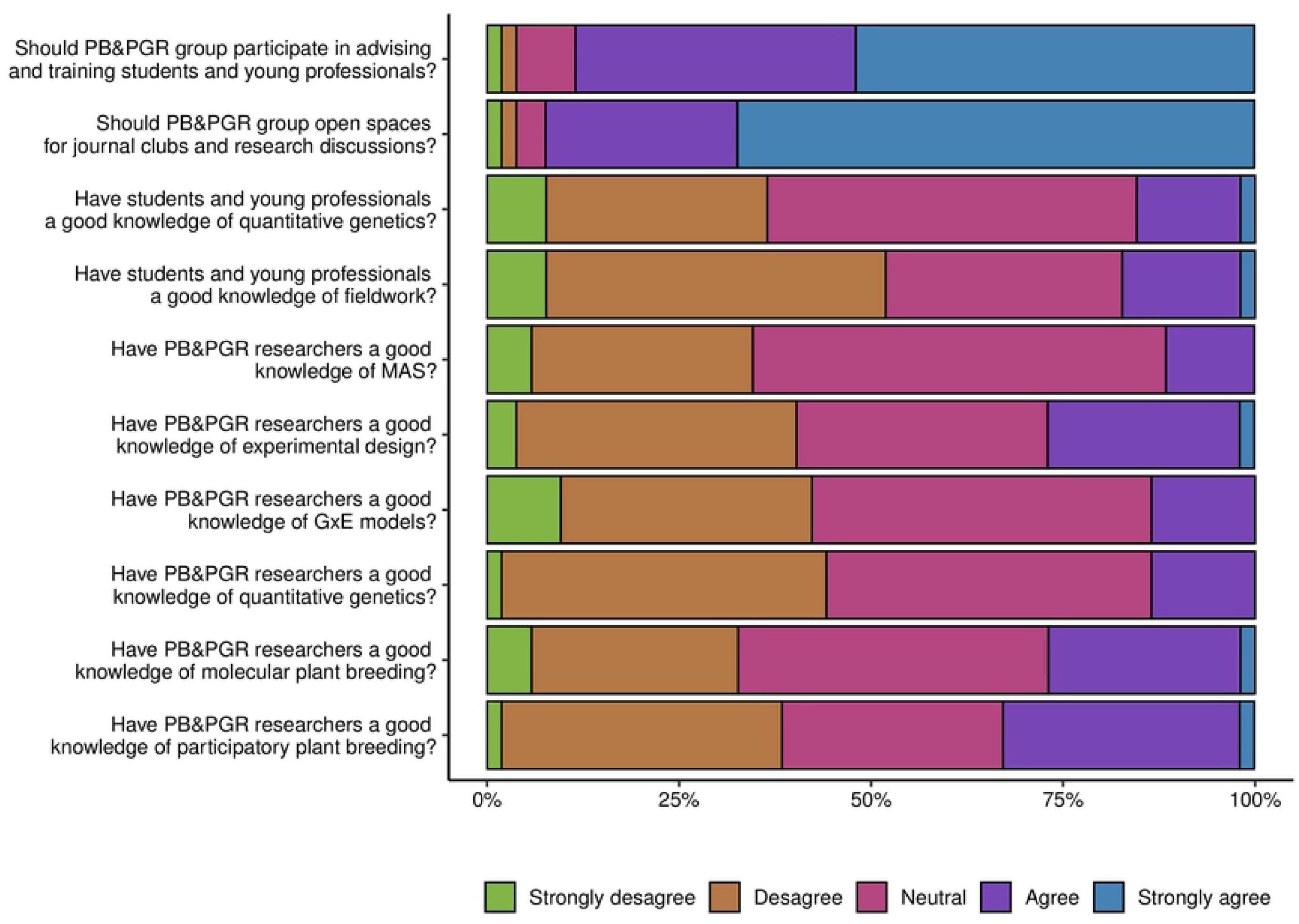
List of ten questions on challenges and opportunities associated with research in plant breeding and plant genetic resources within Agrosavia. The color bars represent the opinion of the 52 researchers surveyed, indicating in percentage the level of agreement for each question. The categories from left to right: strongly disagree, disagree, neutral, agree, strongly agree.

**Figure 6.**
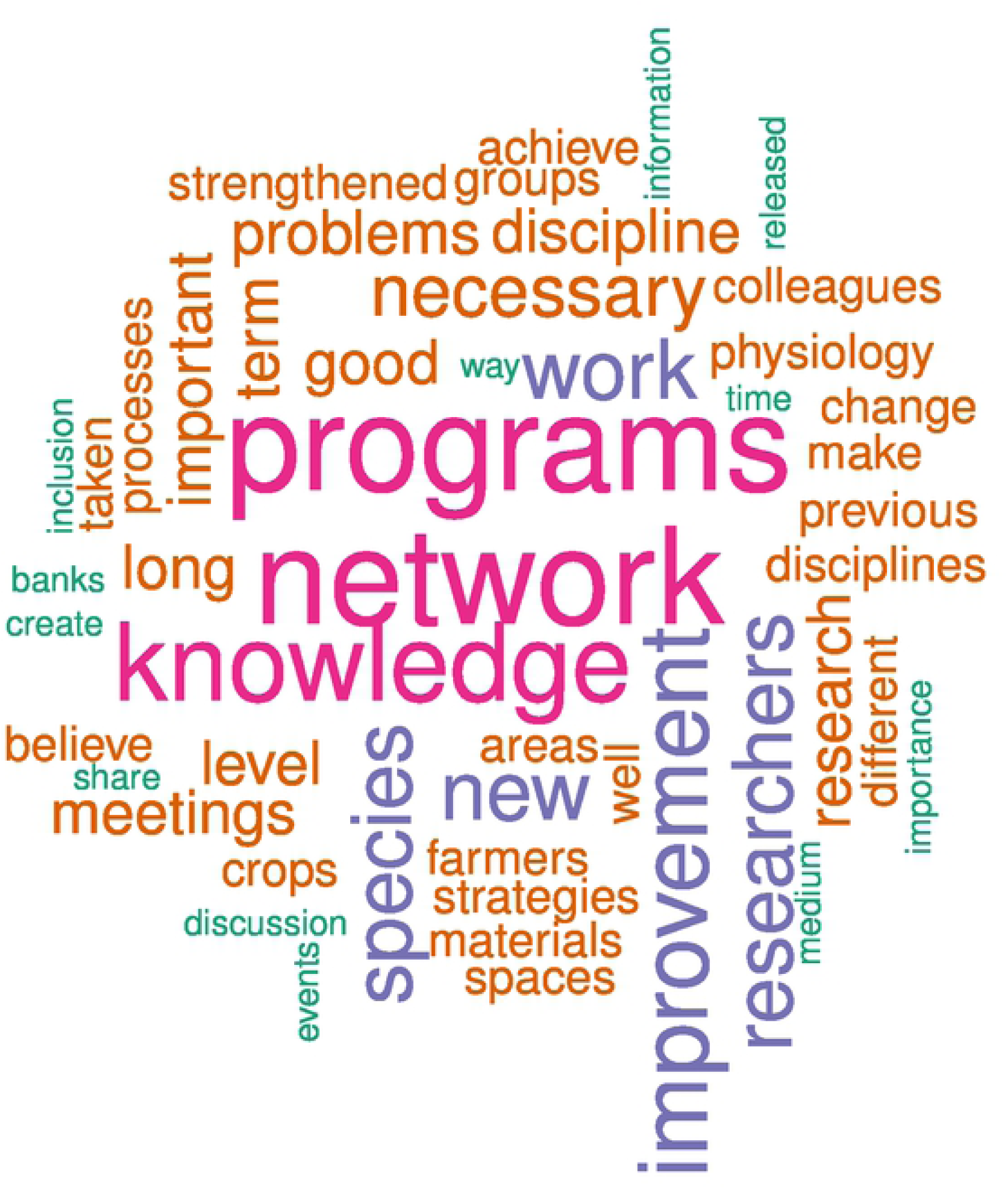
Graphic representation of a word cloud analysis of the 52 opinions about how to improve the plant breeding and plant genetic resources group in Agrosavia.

### Social network

The network had 166 researchers, 48 from the PB&PGR group (with metadata associated) and 118 external, either from Agrosavia or other institutions. The 166 researchers had 812 two-ways interactions among them. We found apparent differences in the number of interactions depending on the types analyzed. The registration of new varieties had the lowest number interaction (N=50), whereas the collaboration in projects in the area had the highest number of interactions (N=242) within the network. Moreover, only two types of links among researchers (i.e., germplasm interchange or request and discussion about new advances in the area) showed more interactions within the PB&PGR group than between PB&PGR and external researchers (Fig. 7).

**Figure 7.**
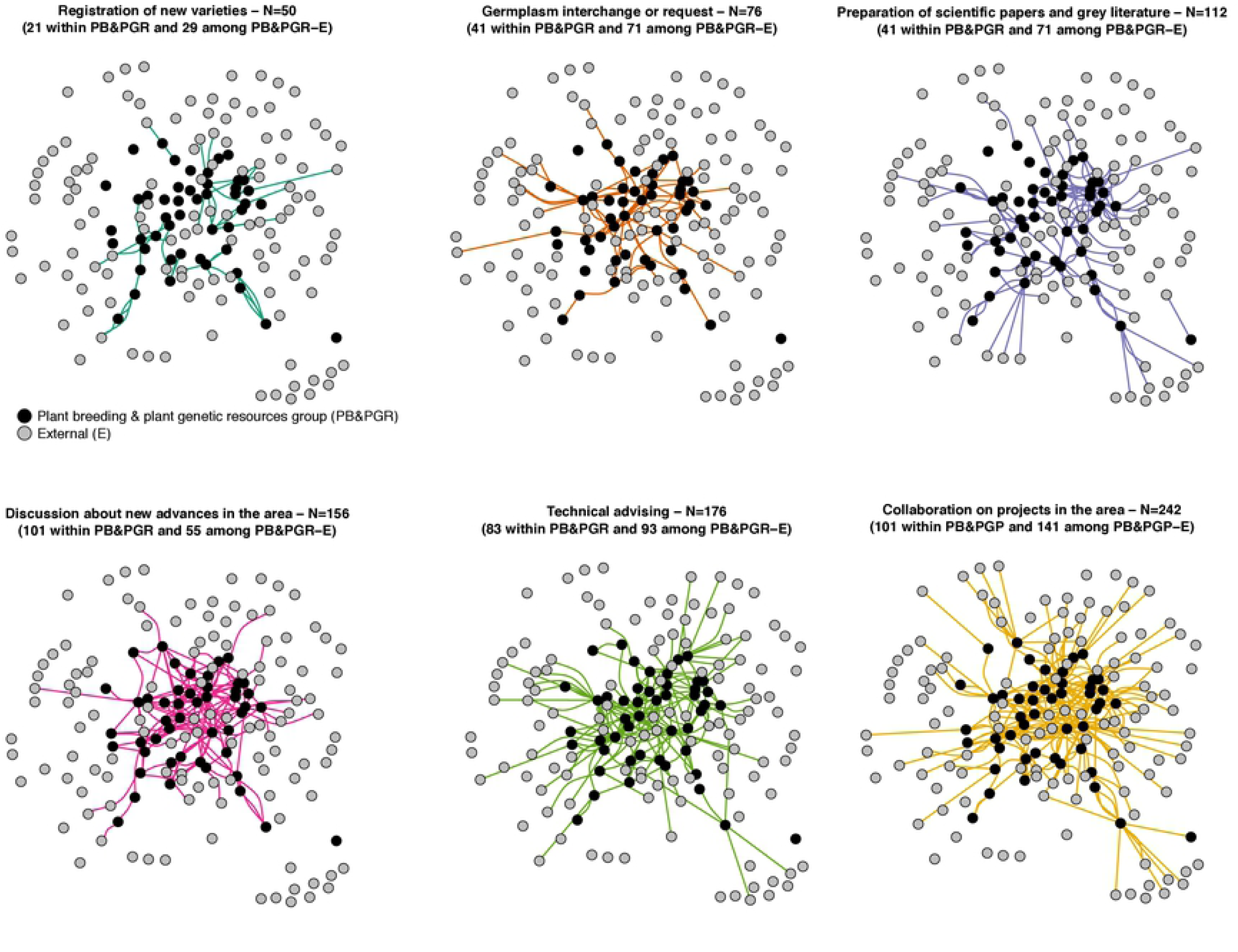
The six types of interactions among 48 researchers from the plant breeding and plant genetic resources (PB&PGR) group in black and 116 external (E), including other disciplines within Agrosavia and outside institutions, in grey. The six types of interactions appear from the lower to the highest number of total connections. Within the parenthesis is specified the number of interactions for both within the PB&PGR group and between the PB&PGR and externals. The dots represent the researchers (i.e., vertex in the network), and the lines represent the interactions (i.e., edges in the system). The dots are in different colors by type from the lower to the highest number of total connections.

Across the network, the diameter was 8, the edge density was 0.03, and the mean distance was 3.07. These values suggest a loose network with many intermediate researchers across interactions among researchers. Likewise, the transitivity was 0.17. This low value indicates a weak system characterized by several discrete and identifiable subgroups.

In the center of the network are the researchers with more influence (i.e., more interactions with other researchers). In contrast, in the periphery are mostly external researchers (Fig. 8A). The top-ten researchers with more interactions (i.e., more degrees in the network) are all from the PG&PGR group, mostly males (N=8) with an M.Sc. degree (N=6) with most of 20 years of experience in the field (N=7) who are currently in the process of retirement during the next two years (Fig. 8B). From them, seven and six researchers are also in the list of the top-ten of hubs and authorities, respectively.

**Figure 8.**
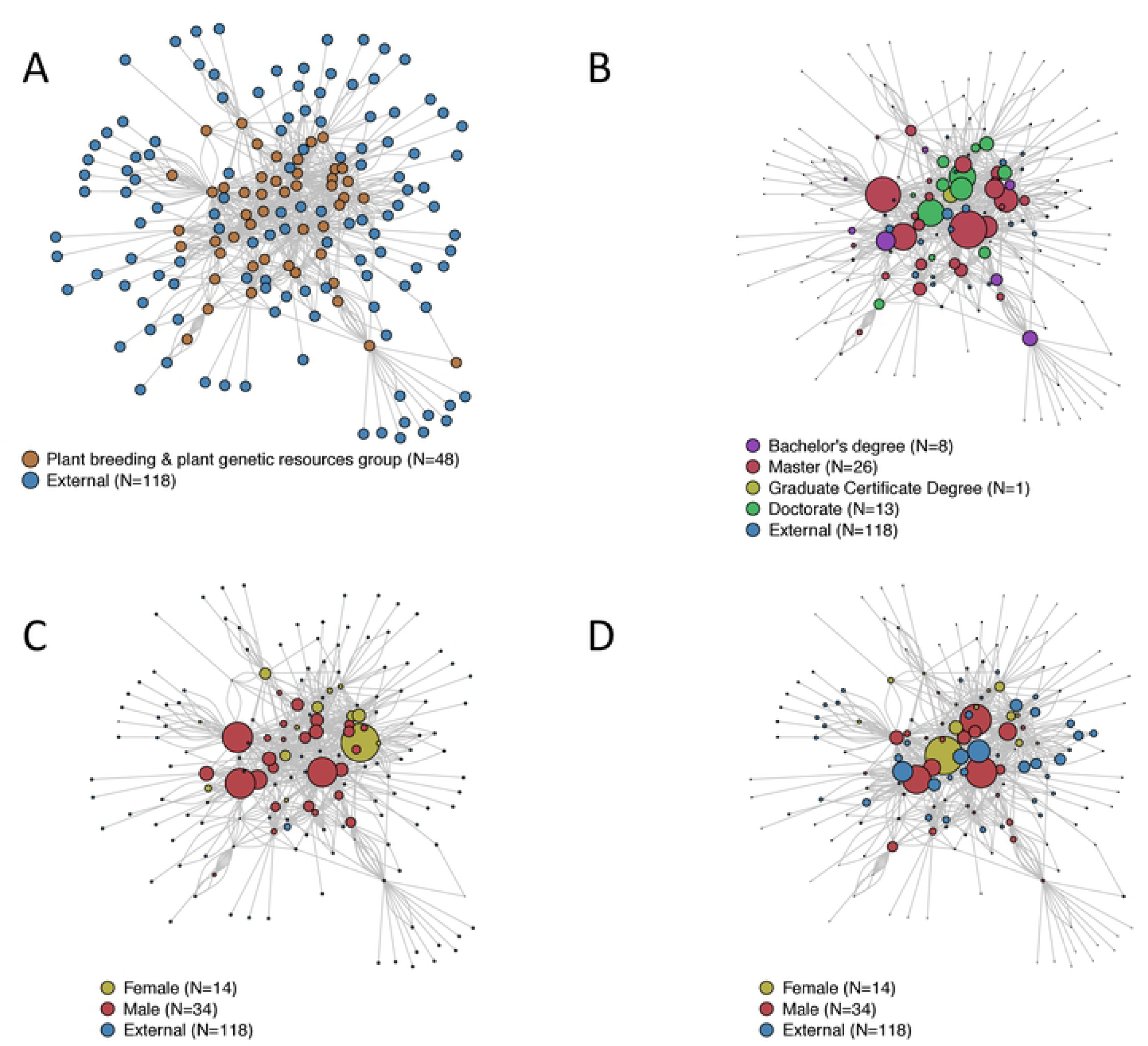
Graphic representation of the network analysis of 48 researchers working in plant breeding and plant genetic resources in Agrosavia and 118 externals, including researchers from other disciplines within Agrosavia and outside institutions. (A) Distribution of researchers (nodes) and their interactions (edges) separating the 48 researchers from the PB&PGR group and 118 external. (B) Characterization of researchers (nodes) by the degree score (i.e., size of the circle) and highest academic degree (i.e., color). The degree score indicates the number of adjacent direct interactions with other researchers. (C) Characterization of nodes by the hub score (i.e., size of the circle) and gender (i.e., color). The hub score indicates the number of outward links that have each researcher with others. (D) Characterization of nodes by the authorities score (i.e., size of the circle) and gender (color). The authority score indicates the number of interactions that receive each researcher from others.

The top-ten of influencer researchers are more diverse (i.e., a broader range of years of experience in the area, and three external researchers in the authorities list). However, similar to the degree score, they are mostly men (i.e., N=8 in both groups) typically with an M.Sc degree (i.e., eigh and six for hubs and authorities, respectively). Spite of females are a minority as influencers; a female researcher got the top score in both hubs and authorities top-ten researchers (Fig. 8C and 8D).

## Discussion

The growing world population, climate challenges, and the loss of agrobiodiversity are the main focus of plant breeders and plant genetic resources managers nowadays. These challenges require that researchers work together using a trans-disciplinary approach to produce results that finally generate an impact in our societies. This study represents the challenges and opportunities that Agrosavia should focus on improving in the next years.

### The population of researchers and their background and collaboration networks

Agrosavia researchers that work in PB&PGR have a high level of scholarly with broad expertise in plant breeding, plant physiology, and plant production, among others. Although this experience is essential for helping move forward the development of a productive research program over the next years in Agrosavia, this study shows the value of decreasing gender gaps, strengthening the generational shift, and improving internal collaboration among different age groups.

Our results showed strong gender bias in the composition of researchers working on plant breeding and plant genetic resources. Although in Agrosavia, the global proportion of male/female researchers was 1.63 in both 2017 and 2018, in our survey, the ratio of 2.45 is worst [15]. Several studies focused on gender equality support that the agricultural sciences are a male dominate research area [16,17]. Consequently, crop sciences, horticulture, and agricultural engineering worldwide describe a female/male global ratio of 0.435, 0.449, and 0.241, respectively [18]. Moreover, according to UNESCO, females working in agricultural and veterinary sciences are just 37.6% of the research population [19].

This gender disparity also agrees with the proportion of male/female ratio graduates in the Agronomy and Veterinary sciences in Colombia. Currently, the ratio is 1.30 for bachelors, 1.27 for M.Sc., and 1.44 for Ph.D. [20]. Moreover, the Agronomy Faculty from the *Universidad Nacional de Colombia* in Bogotá, which represents the largest public university in the country, has a proportion of male/female students of 1.76 and 2.57 among professors staff [21]. For this reason, although this faculty represents only 2.7% of all the student population, it is the third most male-biased across all faculties in the university [22].

Also, our results showed that fewer women researchers in Agrosavia obtained their M.Sc. and Ph.D. degrees abroad compared to men. This result suggests that in Colombia, it is still socially more challenging for women to study overseas compared to men. Unpaid care and housework are strongly biased toward female population in Colombia in two indicators, the total number (i.e., 89.5% of females vs. 62% of males assuming those responsibilities) and time (i.e., 7:03 h for females vs. 3:30 h for male spend for those unpaid tasks in 24h). Thus 12.7% of females declared no free time compared with 8.1% of males [23]. Moreover, female Colombian scientists reported a robust androcentric environment at all levels. Therefore, the combination of unpaid care and housework strongly biased toward women, a patriarchal academic background, and vertical and horizontal segregation pursuing a scientific career may be hindering the mobility of women researchers abroad ([20,22].

On the other hand, the network analysis allowed us to identify that the top-ten of influencer researchers in the area of plant breeding and plant genetic resources within Agrosavia are not evenly distributed neither by the highest degree or the gender, as we hypothesized [24]. The survey showed that the current proportion of researchers with M.Sc. and Ph.D. degrees has a ratio of 2:1. Nine of the M.Sc. researchers have outstanding productivity that has to lead them to reach the category of Associate or Senior. Accordingly, the network showed a concentration of interactions (i.e., degree score) based mainly on male researchers with an M.Sc. degree and with plenty of experience in the area. Likewise, these researchers are the ones who show the highest number of connections with other external actors, probably because their investigative journey has allowed them to expand the number of colleagues with whom they interact. Despite this, the research community is not well consolidated yet. Direct collaboration is scarce, and therefore, it was evident the existence of loosely connected clusters with rare collaborations outside of these subgroups. (Fig. 8). Although Agrosavia needs to focus on these key actors (nodes) as the best way to communicate and transfer innovation within the entire PB&PGR network, the main obstacle is that these identified leaders are close to their retirement age in the next four years.

The network analysis also showed that Agrosavia researchers maintain strong links with many external researchers not included in the survey. This result suggests that those foreign actors are influential consultants for the design of a breeding program strategy. Therefore, the collaborative agreements that Agrosavia is constructing with several institutions inside and outside the country must encourage the formulation of projects that allow them to maintain and strengthen these known external collaborations. Finally, this analysis allowed us to identify productive isolated groups of young researchers that are in the periphery of the network, but that in the future should be in the center of the system, leading projects.

The data collected in this study strongly support the urgency to construct a gender diversity policy combined with a generation shift program in the PB&PGR area. Currently, Agrosavia is a flourishing national institution for agronomic research. Therefore, it is essential that within a gender diversity policy, encourage more interactions among older researchers, characterized by their leadership and commitment, with the creativity and dynamism of younger researchers. This strategy that supports diversity and inclusion would increase Agrosavia’S reputation, making this an attractive working place, especially for women and gender-diverse researchers within the agronomic area.

Currently, Agrosavia is progressively growing the number of Ph.D. researchers formed in Colombia or abroad in the scientific staff [25]. Moreover, Agrosavia has an equal pay policy for females and males across different research rank categories. Therefore, new hiring calls in the area should emphasize this favorable policy to attract more women and gender-diverse researchers in future work calls to replace the current leaders that are close to retirement. Moreover, a gender policy that starts and grows internally could provide an excellent opportunity to promote agricultural development with a gender perspective across different projects lead by Agrosavia [26,27].

Finally, the analysis of improvement networks in corn and wheat in Mexico showed similar patterns that we found in this study [28]. Therefore, this approach could be used in the future for the plant breeding group to manage resistance genes in different crops [29], to understand the dynamics of seed distributions [30] and to determine strategies in grain production [31]. Furthermore, additional analysis using a national survey and publication databases will provide a broad picture of how plant scientists in Colombia are collaborating and what can be improved on institution governance to increase and support these collaborative relationships [32].

### Research experience and skills in the discipline

Researchers working on plant breeding and plant genetic resources in Agrosavia have two noticeable interests for current and future research: 1) tropical fruits for international markets and 2) food sovereignty and food security (Fig. 2). However, results also show that there is not a direct relationship between experience and interest on a specific crop with the impact of new varieties released. For instance, maize was the first crop in the list with a high number of researchers involved in breeding and genetics, but also, with increasing interest to work with in the future. This result makes sense because Colombia consumes 6.2 million tons of maize per year, and this crop is of critical importance for food security and the economy of small farmers. However, Colombia imports more than 70% of the maize mainly from the United States and Argentina, and there is not a public maize breeding program focused on open-pollinated varieties or hybrids to fill the needs of small farmers [33]. Although Mexico faced the same problem as Colombia today, where imports jeopardized the local maize production, they found a solution. In 2010, the CIMMYT (International Maize and Wheat Improvement Center) and SAGARPA (*Secretaría de Agricultura, Ganadería, Desarrollo Rural, Pesca y Alimentación*) of the Mexican Government, successfully started MasAgro. This joint initiative goal is to increase maize productivity, profitability, and sustainability [34]. The MasAgro effort achieved a significant increase in maize production and food self-sufficiency within the country by implementing programs for specific management practices, hybrid seeds, and direct sale in maize markets [35]. Based on this effective result, we suggest constructing similar long-term initiatives and synergies with the Colombian government that support the local production for food security crops, such as maize, beans, quinoa, cassava, and potato, independent of the market trends.

Successful breeding programs worldwide usually start with the use of the broad genetic germplasm base. Thus, collecting, conserving, and characterizing genetic resources are mandatory to introgress novel alleles into elite materials. Colombia has a large National Plant Germplasm Bank with an extensive collection of native and introduced crops [36]. Three current factors are helping improve this situation in the future. First, since 2015, the National Plant Germplasm Bank is working on an ambitious five-year project that aims at implementing a user-friendly GrinGlobal platform for curators and users [37]. Second, as part of the same project, new approaches such as genomics for a robust characterization of collections are beginning to be used [38]. Currently, results for two critical crop collections, such as potato [39]and cacao [40], are available for the public. Soon, we expected to publish similar results for avocado and other native crops. Third, starting in 2018, Agrosavia was officially delegated by the Ministry of Agriculture to manage the Germplasm Banks for food and agriculture [7]. Therefore, we expect that in the next future, a combination of active management, genomic characterization, high-throughput phenotyping, and genomic selection will increase the introgression accuracy of novel germplasm into a breeding program [41,42].

This study also revealed that the base of strategic alliances that are being constructed by Agrosavia with national or regional institutions focused on conventional crops (Fig. 4). For instance, Colombia is validating technologies or varieties generated on other institutions, such as cassava, beans, forages, and rice from CIAT, maize from CIMMYT, and potato from CIP. Further, a new collaboration with high ranked research institutions and universities such as Wageningen University, CIRAD, and USDA are opening new possibilities to co-lead international research initiatives using cutting-edge technologies and novel breeding approaches to switch from being technology adopters to generators of our knowledge and technology.

Besides, the current context of a knowledge-based economy in which experience plays a vital role in economic growth, the growing relationship with universities as a critical player in the national innovation system, becomes essential. An example of this is South Korea that has evolved from being a developing state to a developed country [43]. Thus, the synergistic and collaborative work of Agrosavia with public and private universities has to be a priority. However, the innovation model also involves the industry into the triple helix paradigm where university, industry, and government try to understand and then facilitate and enable collaboration and cooperation between these components to boost national innovation performance [44].

Researchers surveyed expressed worrying issues, such as generational inclusion, powerlessness, capacities, and training opportunities in new areas (Fig. 6 and 7). Our results suggest that researchers are facing a change influenced by the new networking model and the entrance of at least seven new researchers in the area of genetics and plant breeding in the last four years. This situation can explain individual resistance attitudes, usually governed by the anxiety that causes sudden changes in what is conventional or traditional, but also by the broad differences in ages, experiences, scientific productivities, and abilities of the current plant breeding group in Agrosavia [45,46]. Although these results require more detailed analyses, the first approximation suggests that Agrosavia needs to quickly address this assortment of concerns to structure and consolidate a robust PB&PGR group, especially in all those areas that are demanding more time or better training.

## Conclusions

This study is a pivotal input to understand the challenges and opportunities for researchers in the areas of PB&PGR in a state institution as Agrosavia. Based on our analysis, we propose five principles to start working within the next years: 1) The urgent implementation of a gender diversity policy in Agrosavia, combined with a generation shift program that allows contracting a new generation of women and gender-diverse researchers. 2) The construction of long-term initiatives and synergies with the Colombian government to support the local production of food security crops independent of market trends. 3) Better collaboration between the National Plant Germplasm Bank and plant breeding researchers. 4) A concerted priority list of species and external institutions to focus the collaborative efforts in plant-breeding research. 5) Better spaces for the ideation and design of projects among researchers, as well as training programs in border areas associated with plant breeding and plant genetic resources management. Furthermore, we also suggest creating a consultative group to include these five principles across the high-impact research proposals in the generation of new cultivars that respond to biotic and abiotic challenges of national agriculture.

## Acknowledgments

We want to thanks all the colleagues that voluntarily accepted to participate filling the survey and actively discussed the results during the second workshop of plant breeding and plant genetic resources celebrated in March 22-24, 2017. Thanks to Jhon Freddy Hernández for designing the google survey and the pilot analysis of the data obtained. Thanks to the bioethics committee of Agrosavia who evaluated the survey methodology and avalated this study. Finally, all our acknowledge of the Ministry of Agriculture and Rural Development of Colombia for its financial support for this study and all the broader research strategy lead by the Agrobiodiversity Department focused on establishing the plant breeding policy for Agrosavia.

## References

1. Byrne PF, Volk GM, Gardner C, Gore MA, Simon PW, Smith S. Sustaining the Future of Plant Breeding: The Critical Role of the USDA-ARS National Plant Germplasm System. Crop Sci. 2018;58: 451–468. doi: 10.2135/cropsci2017.05.0303

2. FAO. How to Feed the World in 2050. Rome, Italy; 2009.

3. Hunter MC, Smith RG, Schipanski ME, Atwood LW, Mortensen DA. Agriculture in 2050: Recalibrating Targets for Sustainable Intensification. Bioscience. 2017;67: 386–391. doi: 10.1093/biosci/bix010

4. Varshney RK, Graner A, Sorrells ME. Genomics-assisted breeding for crop improvement. Trends Plant Sci. 2005;10: 621–630. doi: 10.1016/j.tplants.2005.10.004

5. Varshney RK, Singh VK, Hickey JM, Xun X, Marshall DF, Wang J, et al. Analytical and Decision Support Tools for Genomics-Assisted Breeding. Trends Plant Sci. 2016;21: 354–363. doi: https://doi.org/10.1016/j.tplants.2015.10.018

6. Lobo M. Importancia de los recursos genéticos de la agrobiodiversidad en el desarrollo de sistemas de producción sostenibles. Corpoica Cienc y Tecnol Agropecu. 2009;9: 19–30. doi: doi.org/10.21930/rcta.vol9_num2_art:114

7. Restrepo Ibiza JL, Gómez Badel A. Una propuesta que parecía improbable: La ruta de Corpoica a Agrosavia. Corporación Colombiana de Investigación Agropecuaria (Agrosavia); 2019.

8. Lemon J. Plotrix: a package in the red light district of R. R-News. 2006;6: 8–12.

9. Wickham H. ggplot2. Wiley Interdiscip Rev Comput Stat. 2011;3: 180–185. doi: 10.1002/wics.147

10. Hu M, Liu B. Mining, and summarizing customer reviews. KDD-2004 - Proceedings of the Tenth ACM SIGKDD International Conference on Knowledge Discovery and Data Mining. 2004. doi: 10.1145/1014052.1014073

11. Bing L, Minqing H, Junsheng C. Opinion Observer?: Analyzing and Comparing Opinions on the Web. Proceedings of the 14th international conference on World Wide Web. 2005.

12. Csárdi G, Nepusz T. The igraph software package for complex network research. InterJournal, Complex Syst. 2006;1695: 1–9.

13. Kleinberg JM. Authoritative sources in a hyperlinked environment. J ACM. 1999;46. doi: 10.1145/324133.324140

14. Hicks DJ, Coil DA, Stahmer CG, Eisen JA. Network analysis to evaluate the impact of research funding on research community consolidation. PLoS One. 2019;14. doi: 10.1371/journal.pone.0218273

15. Vásquez Urriago ÁR, Uribe Galvis CP, Zambrano Moreno GS, Ramírez Gómez MM, Rodríguez Borray GA, Santacruz Castro AM, et al. Balance social 2018. 2019 [cited 24 Jun 2020]. Available: https://repository.agrosavia.co/handle/20.500.12324/35024#.XvN8c82qkVw.mendeley

16. Buttel FH, Goldberger JR. Gender and Agricultural Science: Evidence from Two Surveys of Land-Grant Scientists. Rural Sociol. 2002;67: 24–45.

17. Crowe JA, Goldberger JR. University-Industry Relationships in Colleges of Agriculture and Life Sciences: The Role of Women Faculty. Rural Sociol. 2009;74: 498–524.

18. Larivière V, Ni C, Gingras Y, Cronin B, Sugimoto C. Bibliometrics: Global gender disparities in science. Nature. 2013;504: 211–213.

19. Fernández Polcuch E, Brooks LA, Bello A, Deslandes K. Measuring gender equality in science and engineering: the SAGA survey of drivers and barriers to careers in science and engineering. 2017.

20. Franco-Orozco CM, Franco-Orozco B. Women in Academia and Research: An Overview of the Challenges Toward Gender Equality in Colombia and How to Move Forward. Front Astron Sp Sci. 2018;5. doi: 10.3389/fspas.2018.00024

21. Universidad Nacional de Colombia. Estadisticas docentes: Situación, datos y estadísticas de la planta a marzo de 2018. 2018.

22. Quintero OA. La creciente exclusión de las mujeres de la Universidad Nacional de Colombia. Nómadas. 2016; 123–145.

23. DANE. Boletín técnico Encuesta Nacional de Uso del Tiempo (ENUT) 2016-2017. 2018.

24. Scott J. Social Network Analysis?: A handbook. Newbury Park, CA.: Sage Publications; 1994.

25. Acosta O, Celis J. The emergence of doctoral programmes in the Colombian higher education system: Trends and challenges. Prospects. 2014;44: 463–481. doi: 10.1007/s11125-014-9310-5

26. FAO. Developing gender-sensitive value chains – A guiding framework. Rome; 2016.

27. Holman L, Stuart-Fox D, Hauser CE. The gender gap in science: How long until women are equally represented? PLoS Biol. 2018;16. doi: 10.1371/journal.pbio.2004956

28. Salinas G, Saint C, Cameron M, Wenzl P, Hearne S, Singh S, et al. Avances en el análisis de las redes de mejoramiento genético de maíz y trigo en México. XXV Congreso Nacional y V Internacional de Fitogenética. San Luis Potosí, México; 2014.

29. Garrett KA, Andersen KF, Asche F, Bowden RL, Forbes GA, Kulakow PA, et al. Resistance genes in global crop breeding networks. Phytopathology. 2017;107. doi: 10.1094/PHYTO-03-17-0082-FI

30. Tadesse Y, Almekinders CJM, Schulte RPO, Struik PC. Tracing the seed: Seed diffusion of improved potato varieties through farmers’ networks in Chencha, Ethiopia. Exp Agric. 08/03. 2016; 1–16. doi: 10.1017/S001447971600051X

31. Costabile LT, Vendrametto O, de Oliveira Neto GC, Neto MM, Shibuya MK. Social Network Analysis on Grain Production in the Brazilian Scenario. In: Umeda S, Nakano M, Mizuyama H, Hibino N, Kiritsis D, von Cieminski G, editors. Advances in Production Management Systems: Innovative Production Management Towards Sustainable Growth: IFIP WG 57 International Conference, APMS 2015, Tokyo, Japan, September 7-9, 2015, Proceedings, Part I. Cham: Springer International Publishing; 2015. pp. 36–44. doi: 10.1007/978-3-319-22756-6_5

32. Yu S, Bedru HD, Lee I, Xia F. Science of Scientific Team Science: A survey. Comput Sci Rev. 2019;31: 72–83. doi: https://doi.org/10.1016/j.cosrev.2018.12.001

33. FAO. FAOSTAT Statistical databases. Food and Agriculture Organization of the United Nations. Available: http://faostat.fao.org/. 2017.

34. Camacho-Villa TC, Almekinders C, Hellin J, Martinez-Cruz TE, Rendon-Medel R, Guevara-Hernández F, et al. The evolution of the MasAgro hubs: responsiveness and serendipity as drivers of agricultural innovation in a dynamic and heterogeneous context. J Agric Educ Ext. 2016;22: 455–470. doi: 10.1080/1389224X.2016.1227091

35. Donnet I, Becerril I, Black JR, Hellin J. Productivity differences and food security: a metafrontier analysis of rain-fed maize farmers in MasAgro in Mexico. AIMS Agric Food. 2017;2: 129–148. doi: http://dx.doi.org/10.3934/agrfood.2017.2.129

36. Valencia RA, Lobo A. RM, Ligarreto M. GA. Estado del arte de los recursos genéticos vegetales en Colombia: Sistema de Bancos de Germoplasma. Corpoica Cienc y Tecnol Agropecu. 2010;11. doi: 10.21930/rcta.vol11_num1_art:198

37. Postman J, Hummer K, Ayala-Silva T, Bretting P, Franko T, Kinard G, et al. GRIN-GLOBAL: AN INTERNATIONAL PROJECT TO DEVELOP A GLOBAL PLANT GENEBANK INFORMATION MANAGEMENT SYSTEM. Acta Horticulturae. International Society for Horticultural Science (ISHS), Leuven, Belgium; 2010. pp. 49–55. doi: 10.17660/ActaHortic.2010.859.4

38. Wambugu PW, Ndjiondjop M-N, Henry RJ. Role of genomics in promoting the utilization of plant genetic resources in genebanks. Brief Funct Genomics. 2018;17: 198–206. doi: 10.1093/bfgp/ely014

39. Berdugo-Cely J, Sanchez E, Barrero L, Yockteng R. Genetic Diversity and Population Structure in the Colombian Central Collection of potato (Solanum tuberosum L.) Group Andigenum using SNPs markers. 11 th International Plant Molecular Biology Congress. Iguazú falls, Brazil; 2015.

40. Osorio-Guarín JA, Berdugo-Cely J, Coronado RA, Zapata YP, Quintero C, Gallego-Sánchez G, et al. Colombia a source of cacao genetic diversity as revealed by the population structure analysis of germplasm bank of theobroma cacao l. Front Plant Sci. 2017;8. doi: 10.3389/fpls.2017.01994

41. Araus JL, Kefauver SC, Zaman-Allah M, Olsen MS, Cairns JE. Phenotyping: New Crop Breeding Frontier BT - Encyclopedia of Sustainability Science and Technology. In: Meyers RA, editor. New York, NY: Springer New York; 2018. pp. 1–11. doi: 10.1007/978-1-4939-2493-6_1036-1

42. Yu X, Li X, Guo T, Zhu C, Wu Y, Mitchell SE, et al. Genomic prediction contributing to a promising global strategy to turbocharge gene banks. Nat Plants. 2016;2: 16150. doi: 10.1038/nplants.2016.150 https://www.nature.com/articles/nplants2016150#supplementary-information

43. Lee K-R. University–Industry R&D Collaboration in Korea’s National Innovation System. Sci Technol Soc. 2014;19: 1–25. doi: 10.1177/0971721813514262

44. Jackson P, Mavi RK, Suseno Y, Standing C. University–industry collaboration within the triple helix of innovation: The importance of mutuality. Sci Public Policy. 2017;45: 553–564. doi: 10.1093/scipol/scx083

45. Oreg S, Goldenberg J. Resistance to innovation: Its sources and manifestations. University of Chicago Press; 2015.

46. Rosenberg MJ, Hovland CI. Attitude organization and change: An analysis of consistency among attitude components. Oxford, England: Yale U. Press.; 1966.

